# Heparan sulfate selectively inhibits the collagenase activity of cathepsin K

**DOI:** 10.1101/2024.01.05.574350

**Authors:** Xiaoxiao Zhang, Yin Luo, Huanmeng Hao, Juno M. Krahn, Guowei Su, Robert Dutcher, Yongmei Xu, Jian Liu, Lars C. Pedersen, Ding Xu

## Abstract

Cathepsin K (CtsK) is a cysteine protease with potent collagenase activity. CtsK is highly expressed by bone-resorbing osteoclasts and plays an essential role in bone remodeling. Although CtsK is known to bind heparan sulfate (HS), the structural details of the interaction, and how HS ultimately regulates the biological functions of CtsK, remains largely unknown. In this report, we determined that CtsK preferably binds to larger HS oligosaccharides, such as dodecasaccharides (12mer), and that the12mer can induce monomeric CtsK to form a stable dimer in solution. Interestingly, while HS has no effect on the peptidase activity of CtsK, it greatly inhibits the collagenase activity of CtsK in a manner dependent on sulfation level. By forming a complex with CtsK, HS was able to preserve the full peptidase activity of CtsK for prolonged periods, likely by stabilizing its active conformation. Crystal structures of Ctsk with a bound 12mer, alone and in the presence of the endogenous inhibitor cystatin-C reveal the location of HS binding is remote from the active site. Mutagenesis based on these complex structures identified 6 basic residues of Ctsk that play essential roles in mediating HS-binding. At last, we show that HS 12mers can effectively block osteoclast resorption of bone *in vitro*. Combined, we have shown that HS can function as a multifaceted regulator of CtsK and that HS-based oligosaccharide might be explored as a new class of selective CtsK inhibitor in many diseases that involve exaggerated bone resorption.

## Introduction

Cathepsin K (CtsK) belongs to the papain family of cysteine proteases. The human cysteine cathepsin family includes 11 members and most of them are involved in lysosomal protein degradation(1). Cathepsins can also be secreted to the extracellular space and play active physiological roles at the cell surface and in the extracellular matrix (1). CtsK is unique among cathepsins because it can function as a highly potent collagenase extracellularly (2). The most prominent biological role of CtsK is found in bone resorption. CtsK is highly expressed by osteoclasts, the cells that are solely responsible for bone resorption. Mature osteoclasts actively secrete large quantities of CtsK to degrade type I collagen, the main organic component of the bone matrix (3).

The pivotal role of CtsK in bone homeostasis is manifested by a rare human disease called pycnodysostosis, which is caused by loss-of-function mutations of CtsK (4–6). In addition, deletion of the CtsK gene in mouse resulted in severe osteopetrosis phenotype similar to what is observed in pycnodysostosis (7–11). Due to the essential role of CtsK in bone resorption, CtsK has been pursued as an attractive antiresorptive drug target in treating diseases associated with excessive bone resorption, such as osteoporosis, rheumatoid arthritis and periodontitis (12). One promising CtsK inhibitor, odanacatib, advanced to phase III clinical trials and showed excellent efficacy in reducing fracture risk in postmenopausal women (13, 14). However, the drug was eventually withdrawn due to an increased risk of cardiovascular diseases (12). The unexpected side effect highlighted our incomplete understanding of CtsK biology. One strategy to reduce side effects would be to design CtsK inhibitors that specifically inhibit the collagenase activity while preserving its peptidase activity, which is believed to play essential biological roles in tissues other than bone. In fact, several such inhibitors have been described and molecular modeling studies suggest that they bind to distinct exosites of CtsK, distal from the active site(15, 16).

Like most other proteases, CtsK is synthesized as an enzymatically inactive proenzyme. Activation of CtsK thus requires removal of the N-terminal propeptide by another protease or by CtsK itself. With PIs at 8.4 and 8.7, both proCtsK and CtsK are positively charged proteins and are known to interact with negatively charged glycosaminoglycans (GAGs) (17). The interaction between CtsK and GAGs has been studied since early 2000 (17–19). It appeared that CtsK can bind all major classes of GAGs with various affinities. The interaction between CtsK and chondroitin sulfate (CS) was of particular interest because CS reportedly promotes the collagenase activity of recombinant CtsK expressed in yeast (19, 20). This interaction might have physiological relevance because CS is present in the bone matrix, and has been hypothesized to function as an activator of CtsK for bone matrix degradation. However, the exact physiological role of CtsK-GAG interaction in bone resorption remains unclear and perhaps much more complex because HS, in addition to CS, is also found in the bone matrix and can be expressed by osteoblasts, osteocytes and osteoclasts (21–23). Interestingly, in one study it was found that HS/heparin inhibits the collagenase activity of CtsK, and that the binding affinity of CtsK to HS is much higher than its affinity to CS (17).

To understand the physiological role of CtsK-HS interaction in bone resorption, we examined the structural details of CtsK-HS interaction and how HS might regulate the structure and activity of CtsK. We discovered that HS oligosaccharides, with a length at least 12 sugar residues, can induce mature CtsK to form a stable dimer. Binding of HS to CtsK also greatly increases its structural stability, turning CtsK from a semi-stable enzyme to a highly stable one. We further determined the co-crystal structure of the CtsK-12mer complex, where we found that HS binds to a positively charged surface across multiple Ctsk monomers with a number of the residues confirmed to contribute to strong HS binding. Importantly, we found that HS oligosaccharides can selectively inhibit the collagenase activity of CtsK, and that they can dose-dependently inhibit osteoclast-mediated bone resorption. Our findings strongly suggest that HS functions as an important regulator of the structure and function of CtsK. Deeper understanding of HS-CtsK interactions might lead to novel strategies to treat diseases involving exaggerated bone resorption.

## Results

### CtsK binds HS with high affinity and undergoes HS-dependent dimerization

By heparin-Sepharose chromatography, we determined that 293 cell expressed murine proCtsK binds well to heparin column and elution of proCtsK from the column requires a NaCl concentration of 600 mM NaCl (Fig. 1A). After proCtsK was autoactivated into mature CtsK, we found that mature CtsK displays stronger binding to heparin—requiring 1.4M NaCl for elution from the heparin-column. By surface plasmon resonance analysis, we determined that mature CtsK indeed displays high affinity binding to immobilized HS dodecasaccharide with a *K*_D_ of 6.7 nM (Fig. 1B).

**Figure 1.**
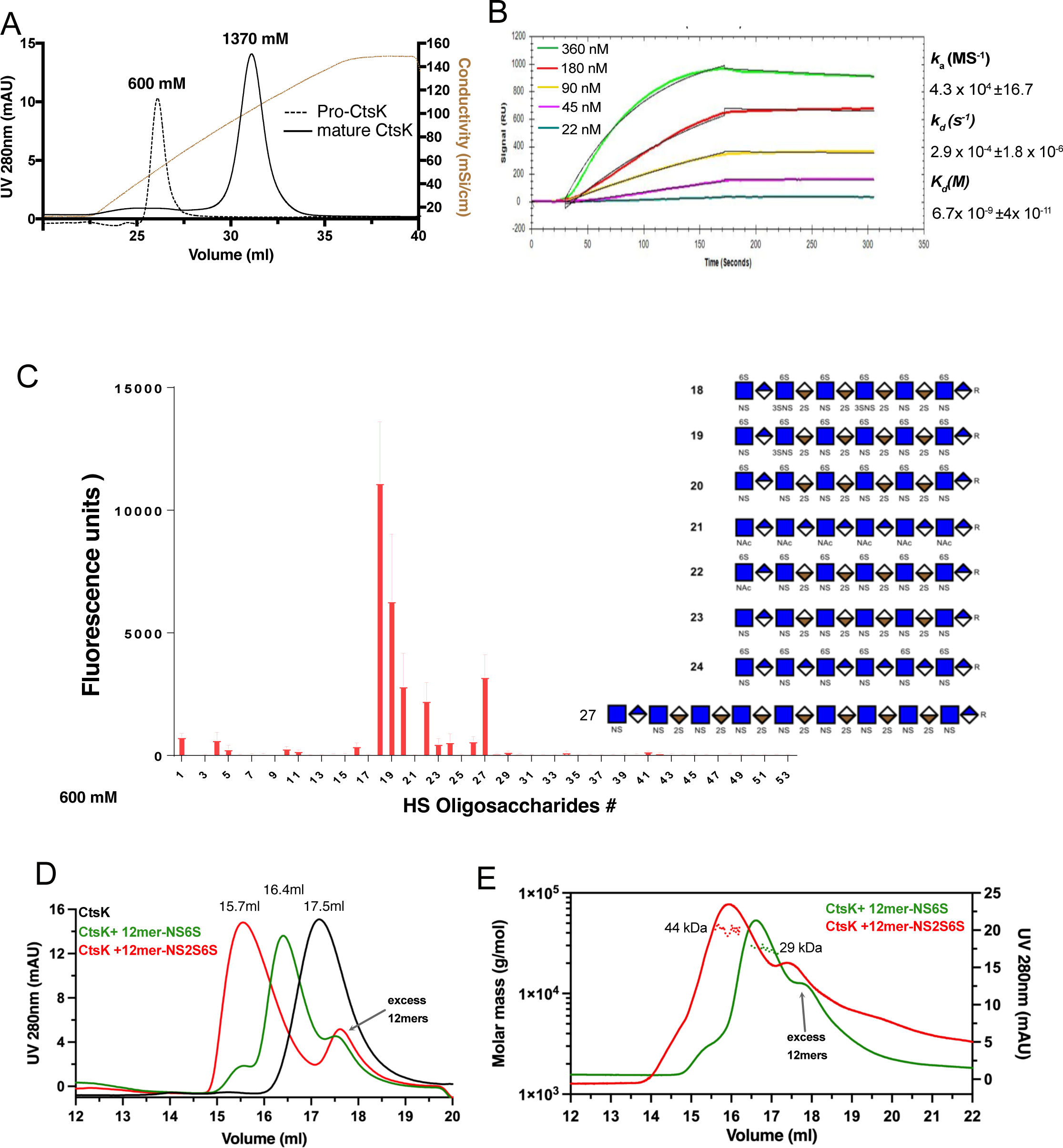
CtsK binds HS with high affinity and undergoes HS-induced tetramerization. Binding of pro-CstK and mature CtsK to heparin-Sepharose column. The brown line represents the salt gradient (150 mM to 2M, measured by conductivity, mS/cm). (B) SPR analysis of binding between mature CtsK and HS dodecasaccharide (HS 12mer). (C) Microarray analysis of mature CtsK binding to HS oligosaccharide microarray. The structure of selected HS oligosaccharides are shown. (D) SEC analysis of CtsK in complex with HS 12merNS2S6S (compound #20) and 12merNS6S (compound #24) on Superdex200 column. (E) MALS-SEC analysis of the MW of CtsK-12mer complexes. The MW data was plotted as dotted lines (left Y-axis) and the UV absorbance was plotted as solid lines (right Y-axis).

To determine the structural specificity of HS-CtsK interaction, we examined binding pattern of CtsK to a HS microarray, which contains 52 structurally defined HS oligosaccharides with various lengths, sequence contexts and sulfation patterns (Figure S1). Interestingly, only a limited number of large HS oligosaccharides, including dodecasaccharides (12mer) and octadecasaccharide (18mer), bind well to CtsK (Fig. 1C). Among different 12mers, we found that under-sulfated 12mers (compound #23 and #24) bind more poorly to CtsK, compared to highly sulfated 12mer (compound #20 and #22). We also observed that 3-*O*-sulfation likely plays a special role in enhancing the CtsK-HS interaction, since compounds #19 and #18 (with one and two additional 3-O-sulfates than compound #20, respectively), display progressively stronger binding to CtsK compared to compound #20. Lastly, we found that increasing the oligosaccharide length can compensate for the deficiency of sulfation, as under sulfated 18mer (compound #27) can bind as well as the highly sulfated 12mer (compound #20 and #22).

Next, we examined whether CtsK-HS interaction could alter the oligomeric state of CtsK, by size exclusion chromatography (SEC) combined with multi-angle light scattering (MALS) analysis. On a Superdex200 SEC column we found that, compared to that of apo CtsK (retention volume 17.5 ml), the retention volume of the CtsK/12mer-NS2S6S (compound # 20, MW = 3.4 kDa) complex was decreased (retention volume 15.7 ml), suggesting that 12mer was able to induce CtsK to form an oligomer.

Interestingly, we also found that when CtsK was mixed with under-sulfated 12mer-NS6S (compound #24, MW = 3.1 kDa), the complex was also eluted at an earlier position (16.4ml) compared to apo CtsK (Fig. 1D). By SEC-MALS analysis we determined that the CtsK/12mer-NS2S6S complex has an estimated MW of 44 kDa, which is very close to the theoretical MW of a CtsK dimer with one molecule of bound 12mer-NS2S6S (49 kDa) (Fig. 1E). Consistent with its later retention position, the MW of CtsK/12mer-NS6S complex is smaller with an estimated MW of 29 kDa, indicating that this is likely a 1:1 complex of CtsK and 12merNS6S (combined theoretical MW = 26 kDa). This result suggests that depending on the sulfation level, HS oligosaccharides can form different oligomeric complexes with CtsK.

### HS oligosaccharides specifically inhibit the collagenase activity of CtsK

To determine how complex formation with HS might alter the enzymatic activity of CtsK, we first examined the peptidase activity of CtsK using a fluorescence-conjugated peptide substrate z-Leu-Arg-AMC. We found that the presence of either heparin or 12mer had no impact on the peptidase activity towards the substrate (Fig. 2A). We further examined the peptidase activity of CtsK using a native multidomain protein. We found that CtsK rapidly digested the extracellular domain of the receptor for advanced glycation end products (RAGE) into multiple fragments, and this process was not affected by the presence of 12mer (Fig. 2B). These results suggest that complex formation between CtsK and HS had no impact on the peptidase activity of CtsK.

**Figure 2.**
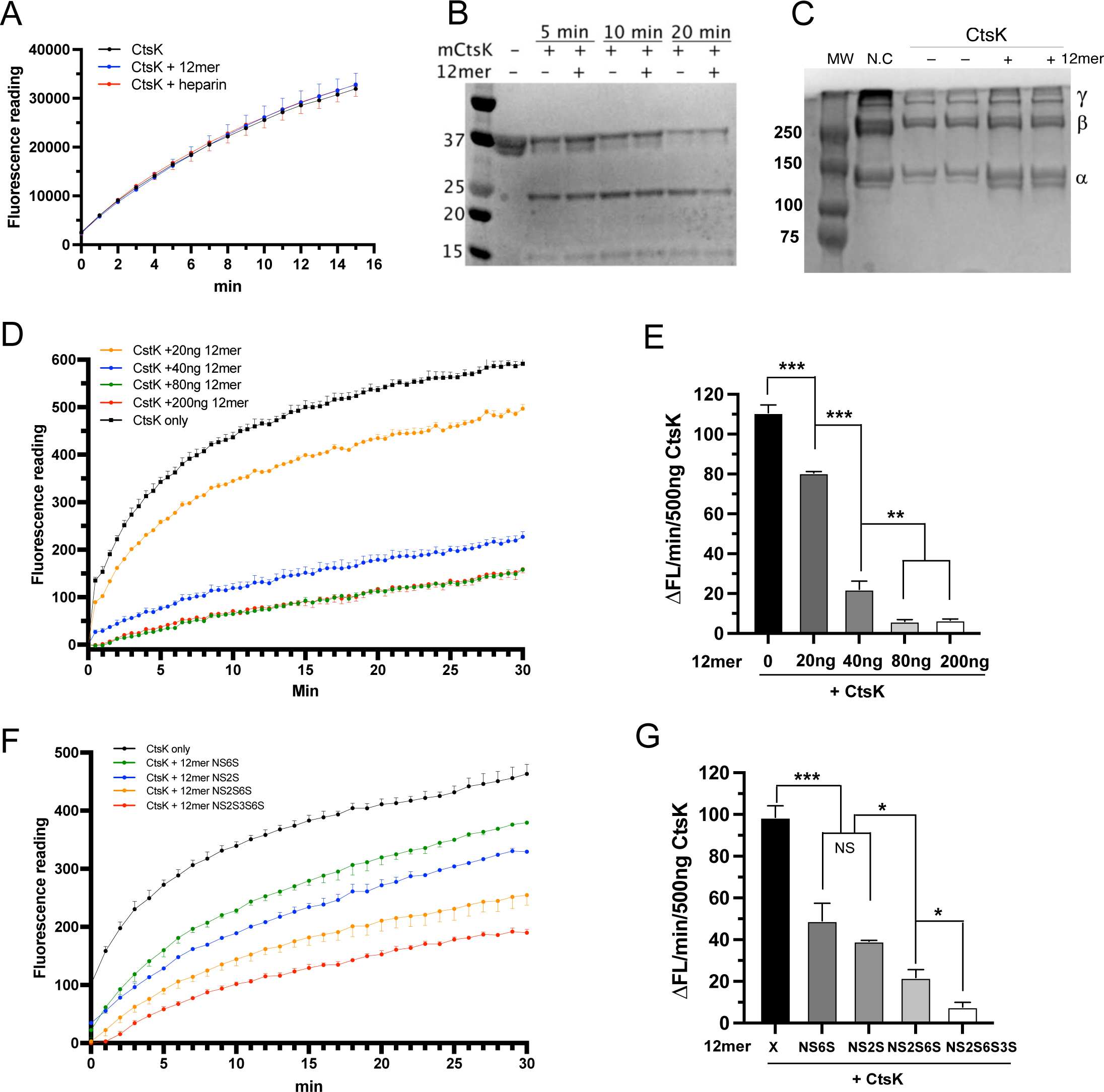
HS selectively inhibits the collagenase activity of CtsK. (A) Digestion of peptide substrate z-Leu-Arg-AMC with CtsK (200 ng) in the absence or presence of HS 12mer (100 ng) and heparin (100 ng). The 12mer used in panels A through E was compound #20. (B) Digestion of the extracellular domain of RAGE with CtsK (20 ng) in the absence or presence of HS 12mer (20 ng) (C) Digestion of type I collagen with CtsK (100 ng) in the absence or presence of HS 12mer (20 ng). N.C, negative control, 5 µg type I collagen. The migration positions of alpha chains, beta chains (cross-linking of two alpha chains), and gamma chain (cross-linking of three alpha chains) of type I collagen are marked on the right. (D) Digestion of FITC labeled type I collagen with 500 ng CtsK in the absence of presence of HS 12mer (20–200 ng). (E) Initial reaction rate (first 2 min) of CtsK derived from plot D. (F) Digestion of FITC labeled type I collagen with 500 ng CtsK in the absence of presence of different HS 12mers (40 ng). (G) Initial reaction rate (first 2 min) of CtsK derived from plot F. For (E) and (G), n = 3, *** represents p < 0.001, ** represents p < 0.01, and * represents p < 0.05 by Student’s t test. Data are representative of at least three separate assays.

Next, we analyzed whether the collagenase activity of CtsK is affected by HS. We found that while the majority of type I collagen was digested by CtsK within 2 hours, the presence of 12mer greatly inhibited the digestion (Fig. 2C). To examine the inhibitory kinetics of 12mer on CtsK, we employed a collagenase assay using FITC-conjugated type I collagen as the substrate. In this assay digestion of the FITC-collagen I conjugate will lead to increased fluorescence signal. We found that 12mer-NS2S6S dose-dependently inhibits the collagenase activity of CtsK (Fig. 2D), and it takes only 40 and 80 ng of 12mer (11.4 and 22.8 pmol) to inhibit 80% and 90% of collagenase activity of 500 ng of CtsK (22.8 pmol), respectively (Fig. 2E). This result also suggests that the stoichiometry of the inhibition is likely 1:2, because one molecule of 12mer was sufficient to inhibit the collagenase activity of two molecules of CtsK to 80%. Next, to understand the correlation between sulfation pattern and inhibitory potency, we performed collagenase assay using 12mers with different sulfation patterns (Fig. 2F). Consistent with our microarray data, we found that 12mer-NS2S (compound #23) and 12mer-NS6S (compound #24), two 12mers that display poor binding to CtsK, display weaker inhibition of CtsK compared to 12mer-NS2S6S and 12mer-NS2S6S3S (Fig 2G).

### HS stabilizes the active conformation of mature CtsK

Similar to what was reported previously (24), we found the purified mature CtsK was semi-stable and gradually lost its activity at 4 °C, even in the presence of the reversible active site inhibitor MMTS (Fig. 3A). By day 3, we observed that around 50% of activity was lost. To understand the underlying reason for this activity loss, we performed SDS-PAGE analysis and heparin-Sepharose column analysis of day-0 and day-3 samples. Heparin-Sepharose analysis revealed that in sharp contrast to day-0 CtsK sample, which eluted as a single peak at 15 ml, day-3 CtsK sample eluted as two peaks, one at the 15 ml position and the other at 11 ml position (Fig. 3B). SDS-PAGE analysis of the Heparin column elution fractions collected from the day-3 CtsK sample revealed that both peaks contains intact CtsK (Fig. 3B, gel picture), confirming there was no protein degradation during the storage at 4 °C. This observation suggests that by day 3 a portion of the CtsK bound with much lower affinity to heparin. Since HS-binding sites of proteins often consist of multiple basic resides arranged in specific spatial configurations (25), a significant reduction in heparin binding often indicates a loss of structural integrity, or a loss of native conformation of HS-binding proteins. Indeed, we found that the CtsK eluted at 11 ml had no detectable enzymatic activity, confirming that this portion of CtsK lost its active conformation.

**Figure 3.**
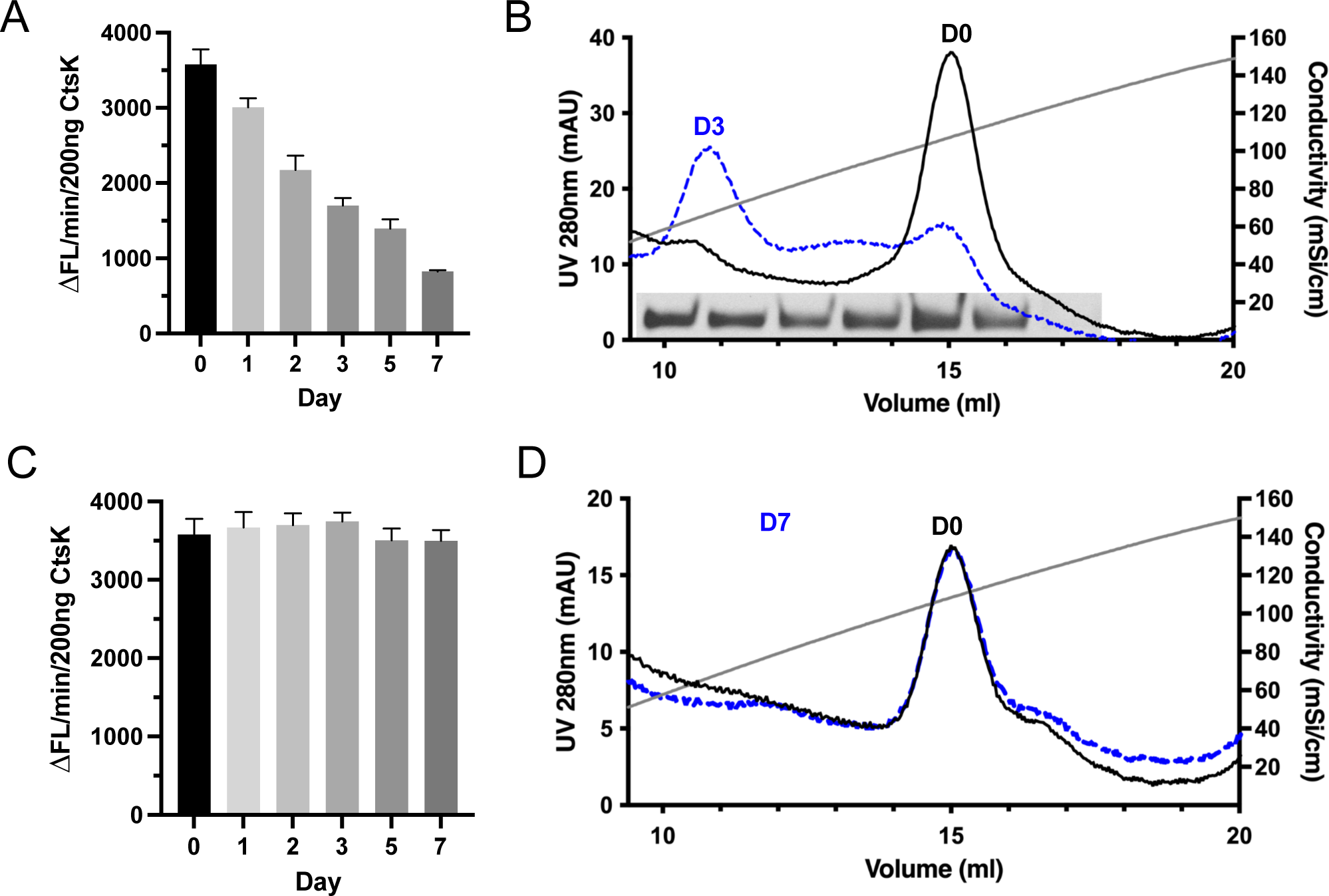
HS stabilizes the active conformation of CtsK. (A) Activity of CtsK after storage at 4 °C in the presence of 5 mM MMTS for up to 7 days. (B) Heparin column elution chromatograms of newly purified mature CtsK before and after storage at 4 °C for 3 days. Coomassie blue staining of Heparin column elution fractions collected from day 3 CtsK samples is shown at the bottom of the chromatogram. Note that all fractions from 10 to 16 ml contain intact CtsK. (C) Activity of CtsK-heparin complex (1:1 molar ratio) after storage at 4 °C for up to 7 days. (D) Heparin column elution chromatograms of CtsK-heparin complex before and after storage at 4 °C for 7 days.

This finding implies there is a connection between the heparin-binding capacity and the active conformation of CtsK, which made us wonder whether we can preserve the active conformation of CtsK using heparin. Indeed, incubation of CtsK with heparin was able to fully preserve the activity of CtsK for at least 7 days (Fig. 3C), and as expected, at day 7, all CtsK that had been incubated with heparin binds to the heparin-Sepharose column at 15 ml position (Fig. 3D). This result suggest that HS might help prolong the half-life of CtsK *in vivo* when they encounter each other and form a complex.

### Co-crystallization of HS 12merNS2S6S with CtsK

To understand how CtsK interacts with HS at the molecular level, we co-crystallized the purified complex of mature CtsK in the presence of a 12mer HS oligosaccharide (12merNS2S6S, compound # 20). These crystals diffract to a resolution of 3.1 Å and contain two molecules of CtsK (molecules A and B) in the asymmetric unit that are related by a non-crystallographic two-fold (Fig. 4A and B). Residues from Lys41-Lys44 and Lys103-Lys106 of each monomer are found at the interface burying 452 Å^2^ in surface area, accounting for 4.7% of the accessible surface of each monomer (Fig. 4B). This arrangement results in few specific interactions between molecules A and B, and is not predicted to be a physiological dimer based on PISA analysis(26). However, this interaction has also been observed in another crystal structure of mouse CtsK (PDB ID code 5t6u) (27). Substantial electron density was found along a two-fold crystallographic axis that could be modeled with an octasaccharide HS with 50% occupancy in each direction across the two-fold axis, interacting with positively charged surfaces from both CtsK mol A and symmetry related mol A’ (Figure S2). Additional density is observed at the termini of the oligosaccharides extending across the face of the B and B’ (symmetry-related monomer) molecules in the crystal lattice. Unlike the monomer A/B interface, there are no clear interactions between the two CtsK symmetry related molecules A/A’ that are bridged by the HS (Fig. 4C). Residues Arg108, Arg111, Lys119, Arg123, Arg127, Lys214 from both A and A’ molecules are in position to form interactions with the visible octasaccharide in either orientation of the HS with Lys176 also located near the binding site. Interestingly, extending the oligosaccharide beyond the octasaccharide suggests residues Arg108, Arg111, Arg 123, Arg127, and Arg214 from molecules B or B’ as well, could potentially participate in HS binding. This arrangement suggests HS could stabilize either a A/A’ dimer with almost no protein-protein interactions or stabilize a A/B dimer with weak protein-protein interactions. Interestingly, the position of HS within this crystal structures differs from that of chondroitin sulfate binding in two different structures (Fig. 4D). In one crystal structure (pdb ID code 4N8W) of CS binding to CtsK, the CS oligosaccharide is found binding perpendicular to that of HS with only partial overlap with the HS-binding site(28). In another crystal structure of CS binding to CtsK (pdb ID code 3C9E), the CS binding site is found distal from that of HS binding (29). These comparisons support the suggestions that HS and CS can interact differently with CtsK and thus have different effects on CtsK function.

**Figure 4.**
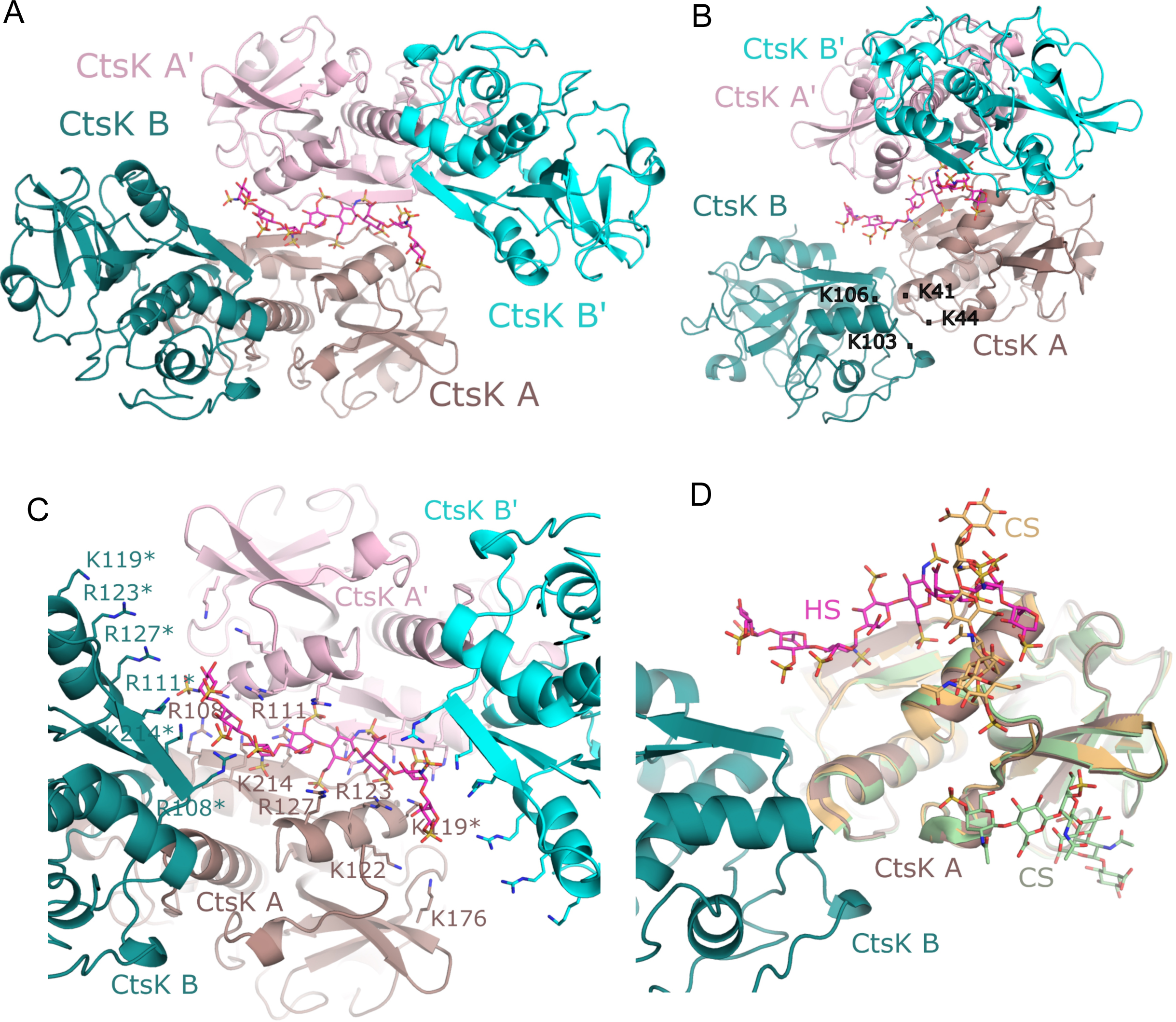
Crystal Structure of CtsK-HS complex. (A) Crystal structure of CtsK with 12merNS2S6S bound. The asymmetric unit is composed of CtsK monomer A (dark salmon), CtsK monomer B (dark teal), and HS 12merNS2S6S (magenta). A crystallographic symmetry related unit is displayed in pink (monomer A’) and cyan (monomer B’). (B) Rotation of figure A displaying the small interface between CtsK monomers A and B and the extend of HS along the surface of monomer B. (C) Binding of HS to CtsK. Displayed are the residues found along the HS binding path. Residues from CtsK monomers A and B are labeled while those from A’ and B’ are shown without label. Symmetry related HS has been omitted for clarity. Asterisks denotes residues for which sidechains were modeled for this figure as they lacked clear sidechain density in the structure. (D) Superpositions of crystal structures of human CtsK with bound chondroitin sulfate (CS) (PDB_idcode 4N8W in orange; PDB_idcode 3C9E in light green) onto the asymmetric unit of mCtsK with 12merNS2S6S.

### CtsK-Cystatin C heterodimer

Previously our lab determined that a widely expressed cysteine protease inhibitor, Cystatin C (Cst3) as a HS-binding protein under acid pH (30). The fact that both Cst3 and CtsK bind HS raises the possibility that HS might regulate the inhibitory activity of Cst3 towards CtsK. To test this, we examined the enzymatic activity of CtsK towards a peptide substrate and found that Cst3 potently inhibited the activity of CtsK. However, the presence of 12mer (compound # 19) neither enhanced nor interfered with the inhibitory activity of Cst3 towards CtsK in this assay (Fig. 5A). By SEC, we found that CtsK (22 kDa) and Cst3 (13 kDa) readily form a stable complex, which eluted at a slightly reduced retention volume than the 44 kDa MW standard (Fig. 5B). Based on this retention volume, we predict that the complex has a stoichiometry of 1:1, and the complex has a MW of 35 kDa. Interestingly, when CtsK, Cst3 and 12mer-NS2S6S were co-incubated, they formed a larger complex with a retention position very close to 66 kDa. This retention position suggests that the ternary complex most likely includes two molecules of CtsK and two molecules of Cst3, which would give a combined MW of 70 kDa.

**Figure 5.**
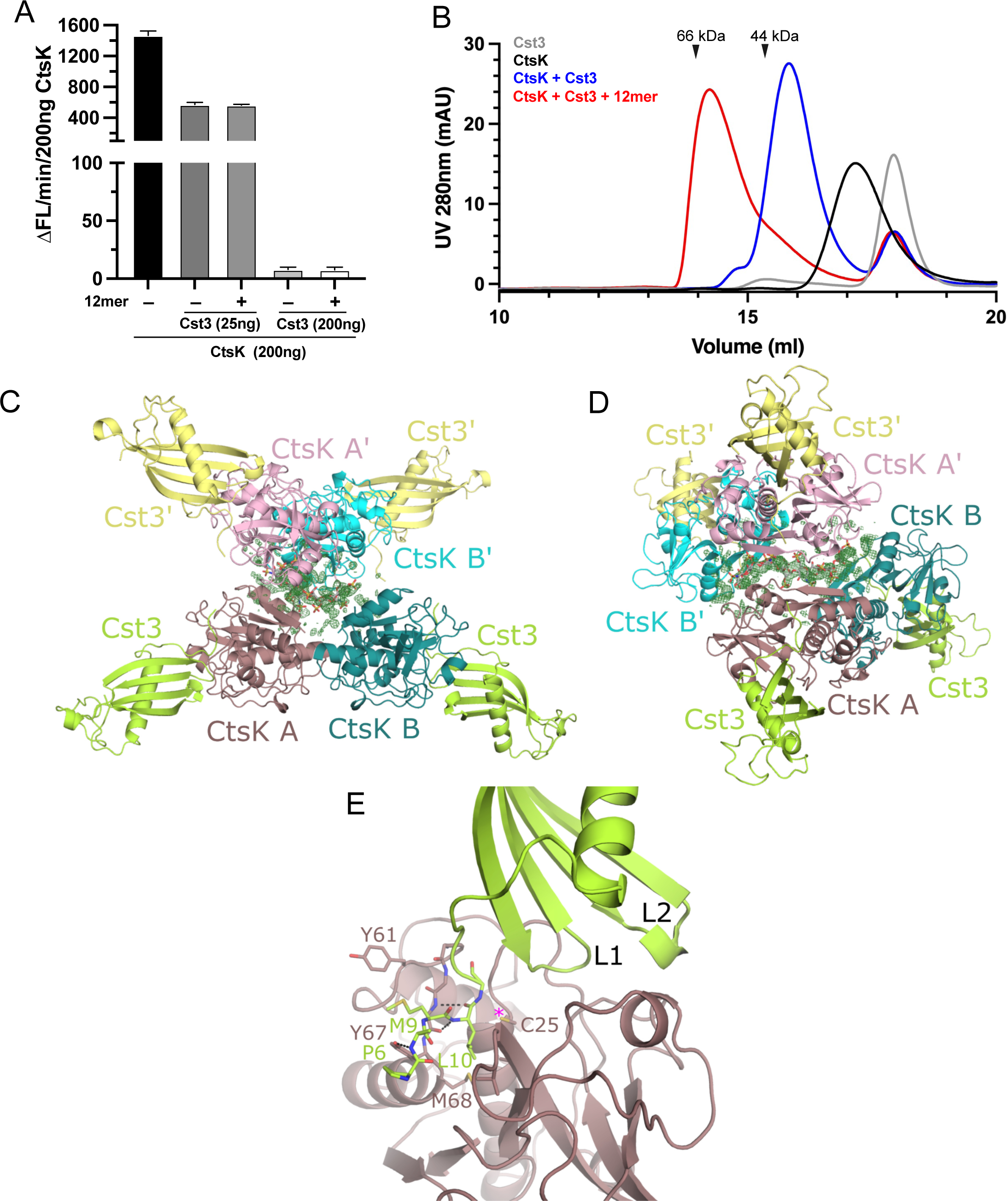
Structure of HS-CtsK-Cst3 ternary complex. (A) Digestion of peptide substrate with CtsK (200 ng) in the absence or presence of Cst3 (25 or 200 ng), with or without HS 12mer (100 ng). (B) SEC analysis of CtsK complex formation with Cts3, in the presence of absense of HS 12merNS2S6S on Superdex200 column. (C) Crystal structure of CtsK bound to Cst3 in the presence of 12merNS2S3S6S. The asymmetric unit is composed of CtsK monomer A (dark salmon), CtsK monomer B (dark teal), two Cst3 molecules (both colored lime). A crystallographic symmetry related unit is displayed in pink (monomer A’), cyan (monomer B’), and two Cst3 monomers (both pale yellow). Difference electron density found at the crystallographic axis is shown in green contoured at 2.0σ. The 12merNS2S6S from the CtsK/12merNS2S6S structure has been superimposed based on CtsK monomer A and is shown (gray) for perspective. (D) alternate perspective of figure C displaying the continuous nature of the difference electron density for HS. (E) Interface of CtsK with Cst3 formed by loops L1 and L2 of Cst3 and the N-terminal residues of Cst3 that bind to the active site of CstK blocking substrate binding and access to the catalytic Cys25 (denoted with pink asterisk).

### Co-crystallization of HS 12mer2S3S6S with CtsK-Cystatin C complex

To understand the architecture of this ternary complex, we solved the crystal structure of the Cst3-Ctsk complex in the presence of 12merNS2S3S6S (compound # 19) oligosaccharide to 2.75 Å. In this structure, the asymmetric unit consists of two complexes of CtsK-Cst3 that form an interaction through a similar CstK/CstK (monomers A and B) interface as seen in the CtsK-12mer crystal structure, involving loops containing residues Lys41-Lys44 and Lys103-Lys106 of each monomer with the Cst3 bound at the active site, distal from the CtsK/CtsK interface in the crystal (Fig. 5C & D). As observed in other cystatin-cathepsin co-crystal structures (31, 32), Cst3 binds CtsK by three structural motifs: the N-terminal coil (Pro7-Ala12), a central β-hairpin loop L1 (Leu56-Gly59) and a C-terminal hairpin loop L2 (Pro105-Lys107) (Fig. 5E). This positions the N-terminal coil and L1 in close proximity to the catalytic cysteine (Cys25) blocking substrate access to the active site. In this structure, clear electron density is seen for N-terminal residues Pro7-Gly11 that form extensive interactions with the active site of CtsK blocking substrate access to Cys25 through both hydrogen bonding and hydrophobic interactions involving residues Met9 and Leu10 (Fig. 5E).

In the presence of Cst3, the complex crystallizes in a different space group than the Ctsk-12merNS2S6S crystal structure, yet electron density is found across a crystallographic axis between two symmetry related molecules of CstK (Fig. 5C & D). forming a similar crystallographic tetrameric arrangement of CtsK as was seen in the binary structure of CtsK in the presence of 12merNS2S6S (Figure S3). Although the density is too weak to model HS into, superposition of CtsK from the CtsK-12merNS2S6S structures onto the CtsK in this complex positions the HS at the same interface between the symmetry related CtsK molecules as well as extending across CtsK molecule B in the asymmetric unit (Fig. 5C & D). Thus, despite the presence of Cst3, HS can form the same interactions with CtsK as seen in the CtsK-12merNS2S6S complex.

### Mutagenesis of the HS-binding site of CtsK

To verify the HS-binding site observed in the co-crystal structures (Fig. 4C), we performed site-directed mutagenesis to determine the relative contributions of the identified HS-binding residues. The HS binding capacity of eight single alanine substitution mutants were first determined by heparin-Sepharose chromatography. Among the eight, we found 4 mutants–R111A, R123A, R127A and K214A–had the most dramatic effect (>200 mM reduction in NaCl elution concentration) on weakening the binding of CtsK to heparin (Table 1). Interestingly, in our co-crystal structure (Fig. 4E) these four residues are centrally located within the HS-binding site and are all capable of making electrostatic interactions with multiple sulfate groups of 12mer. The arginine sidechains in particular, can make multiple interactions simultaneously. In addition, R108A and K176A mutants were also able to weaken heparin-binding to a moderate degree (reduction in NaCl elution concentration of 150 and 90 mM, respectively). The milder effect of these two mutants likely reflects their more peripheral location within the HS-binding cleft and suggests how they could contribute to binding of longer oligosachcarides(Fig. 4C). Unexpectedly, mutation of R119 and R122, two residues that appear not too far away from the bound 12mer (Fig. 4C), had no effect on heparin binding whatsoever. This last finding suggests that despite the presence of large number of basic residues, the HS-binding site of CtsK is quite defined. As expected, combining two (R123A-R127A) or three (R111A-R123A-R127A) of the single mutants resulted in much greater reduction in HS-binding capacity (Table 1).

**Table 1.** CtsK mutants binding to heparin Sepharose column.

| **Mutant** | **Elution NaCl concentration (mM)** | **Reduction compared to WT (mM)** |
| --- | --- | --- |
| WT-CtsK | 1370 | — |
| R108A | 1220 | 150 |
| R111A | 1160 | 210 |
| K119A | 1370 | 0 |
| K122A | 1370 | 0 |
| R123A | 1150 | 220 |
| R127A | 1150 | 220 |
| K176A | 1280 | 90 |
| K214A | 1135 | 235 |
| R123A-R127A | 960 | 410 |
| R111A-R123A-R127A | 880 | 490 |

These HS binding-impaired CtsK mutants can serve as excellent tools to investigate the biological significance of HS-CtsK interaction. Consistent with the greatly reduced binding to heparin-Sepharose (Table 1), we found that the double (R123A-R127A) and triple mutant (R111A-R123A-R127A) both displayed poor binding to MC3T3 cell surface HS (Fig. 6A). Clearly, mutations of HS-binding residues do not affect either the peptidase or the collagenase activity of CtsK, as the triple mutant functions equally well as the WT CtsK (Fig. 6B and C). However, in sharp contrast to WT CtsK, whose collagenase activity was robustly inhibited by 12mer (compound #20) (Fig. 6D and E), the triple mutant is no longer inhibited by 12mer (Fig. 6D and F. Combined, these results strongly suggest that CtsK-HS interaction bears a special significance in regulating the collagenase activity of CtsK, which might impact how CtsK functions during bone resorption.

**Figure 6.**
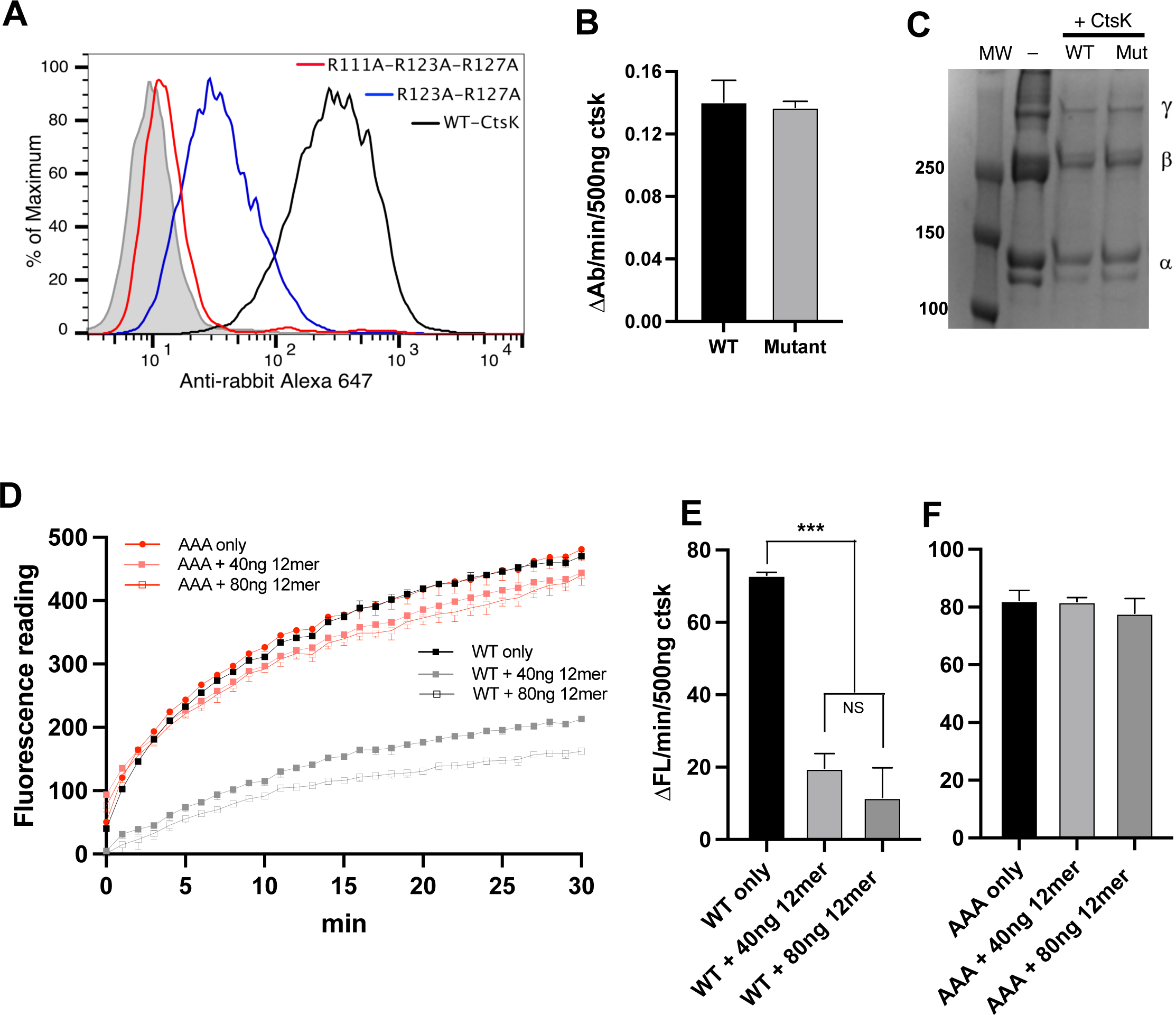
HS-binding deficient CtsK maintains normal collagenase activity but is no longer inhibited by HS. (A) Binding of WT CtsK and CtsK mutants (R123A-R127A and R111A-R123A-R127A) to MC3T3 cell surface was determined by FACS. The bound CtsK was detected by staining with a rabbit anti-CtsK antibody, followed by anti-rabbit-IgG Alexa-647. The shaded histogram is from cells stained only with primary and secondary antibodies. (B) Enzymatic activities of WT CtsK and R111A-R123A-R127A CtsK were measured by a peptide substrate. (C) Digestion of type I collagen with WT CtsK or R111A-R123A-R127A mutant. (D) Digestion of FITC labeled type I collagen with 500 ng WT or mutant CtsK in the presence of absence of HS 12mer (40 or 80 ng). (E) Initial reaction rate (first 2 min) of WT CtsK derived from plot D. (F) Initial reaction rate (first 2 min) of mutant CtsK derived from plot D. n = 3, *** represents p < 0.001, by Student’s t test. Data are representative of at least three separate assays.

### Osteoclasts highly express HS

Our findings so far point to a potential role of CtsK-HS interaction in regulating CtsK activity during bone resorption. To further substantiate this notion, we performed histological analysis of murine femur sections to examine the expression pattern of HS and CtsK in bone tissue. As expected, immunostaining of CtsK showed clearly that osteoclasts highly express CtsK and they are the only cells that express detectable amount of CtsK in bone tissue (Fig. 7A). Immunostaining of HS by a highly specific human anti-HS mAb (HS20) revealed that osteoclasts strongly express HS at a level that is much higher than other HS-expressing cells in bone tissue, such as chondrocytes and osteoblasts (Fig. 7B, indicated by black asterisks or arrows, respectively). In fact, the HS-staining of osteoclasts is so distinct that it can be used as a reliable way of visualizing osteoclasts in bone sections. When bone sections were co-stained with anti-CtsK antibody and HS20, we found CtsK and HS colocalize extensively in osteoclasts (Fig. 7C). This finding strongly supports the hypothesis that that CtsK and HS may interact with each other during osteoclast-mediated bone resorption.

**Figure 7.**
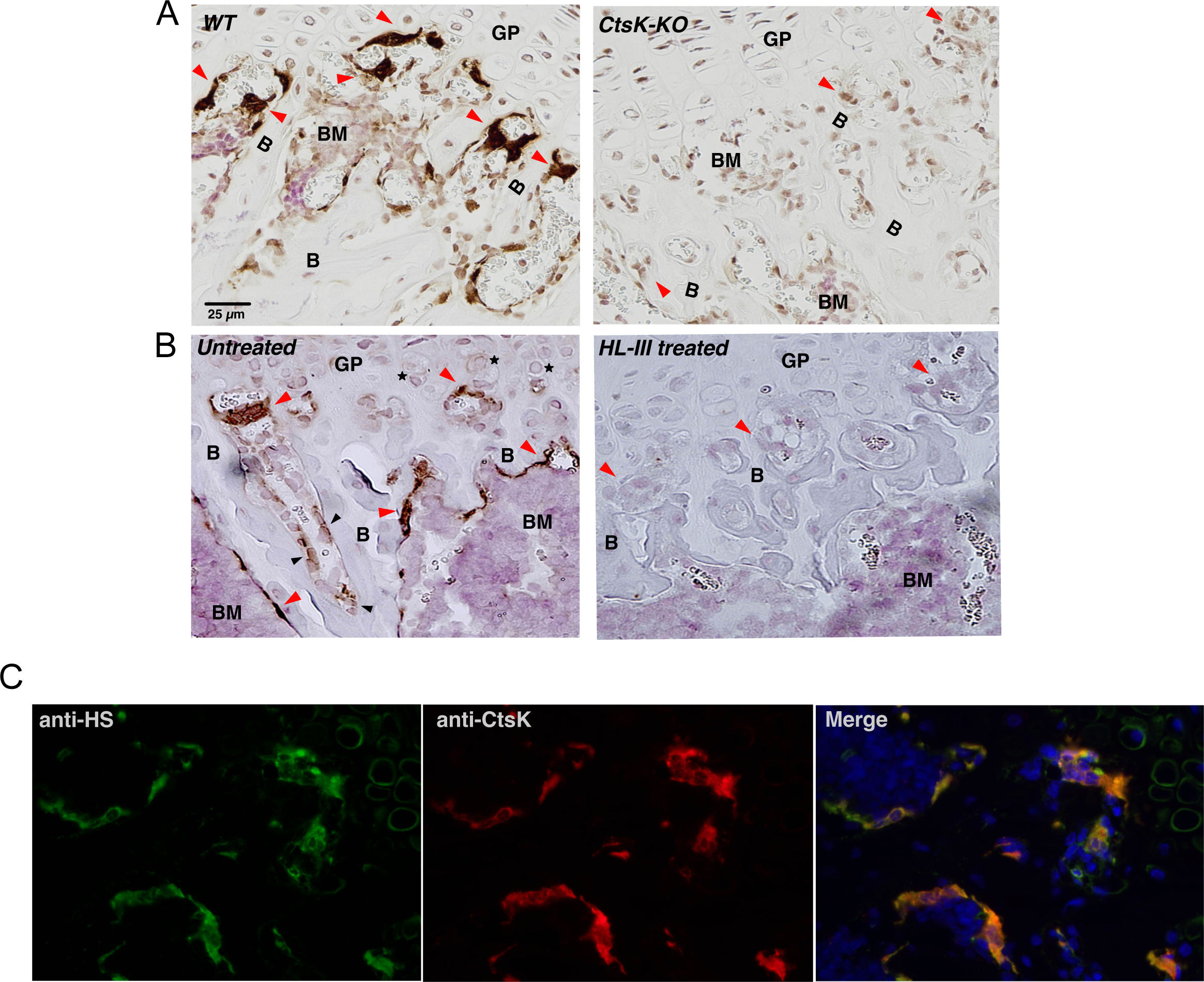
Osteoclasts highly express HS. (A) WT or CtsK-KO mouse femur sections were stained with a rabbit polyclonal anti-CtsK antibody and developed using IHC method. Nuclei were counterstained with Ehrilch’s hematoxylin. osteoclasts are indicated with red arrowheads, which were strongly stained in WT femur, but not in CtsK-KO femur. B, bone; BM, bone marrow; GP, growth plate. Magnification = ×200. (B) WT mouse femur sections, with or without pretreatment of heparin lyase III, were stained with a human anti-HS mAb (HS20) and developed using IHC method. Osteoclasts in untreated slide were strongly stained with HS20. Chondrocytes are indicated by black stars, and osteoblasts are indicated by black arrowheads. (C) Co-localization of CtsK and HS in osteoclasts. WT mouse femur section was co-stained with anti-CtsK antibody and HS20 and visualized with anti-rabbit Alexa594 and anti-human Alexa488 secondary antibodies. Magnification = ×200.

### HS 12mer inhibits bone resorption by osteoclasts

The fact that HS oligosaccharides selectively inhibit the collagenase activity of CtsK suggests that they could potentially be developed into anti-resorption therapeutics. To test the anti-resorption activity of HS oligosaccharides, we employed an *in vitro* bone resorption assay where bone marrow macrophage-derived osteoclasts are cultured on bovine cortical bone slices. Seven days after seeding of osteoclasts onto bone slices, we observed extensive bone resorption pits densely packed on the bone surface (Fig. 8A. positive control panel). However, if 12mer-NS2S6S (compound #20, 25 µg/ml) was added into the culture medium for the last 6 days of bone resorption, we found the total area of bone resorption was inhibited by 75% (Fig. 8A and B). On bone slices treated with 12mer we noticed that while the resorption pits can be easily identified on the bone surface, the pits were much more superficial (lighter blue stain compared to darker blue stain in the positive control, Fig. 8A insets, top two panels), which is consistent with reduced resorption capacity. We further tested the inhibitory activity of 12mer-NS2S6S3S (compound #19), which in our earlier test showed higher anti-collagenase activity compared to 12mer-NS2S6S (compound #20) (Fig. 2F and G). As expected, 12mer-NS2S6S3S also dose-dependently inhibits bone resorption, and the 25 µg/ml dose was able to inhibit more than 90% of bone resorption (Fig. 8A and B). Interestingly, the inhibitory effect of 12mer-NS2S6S3S at 5 µg/ml was very similar to the inhibitory effect of 12mer-NS2S6S at 25 µg/ml (Fig. 8B), which suggests that the inhibitory potency of 12mer-NS2S6S3S is 5 times higher than that of 12mer-NS2S6S. To determine whether treatment with the 12mers had any effect on the attachment and survival of seeded osteoclasts, we performed TRAP staining to visualize osteoclasts at the completion of the treatment. We found that the density and morphology of the 12mer-treated osteoclasts are indistinguishable from untreated osteoclasts (Fig. 8C and D). This result suggests that HS 12mers inhibits bone resorption without affecting the survival of osteoclasts.

**Figure 8.**
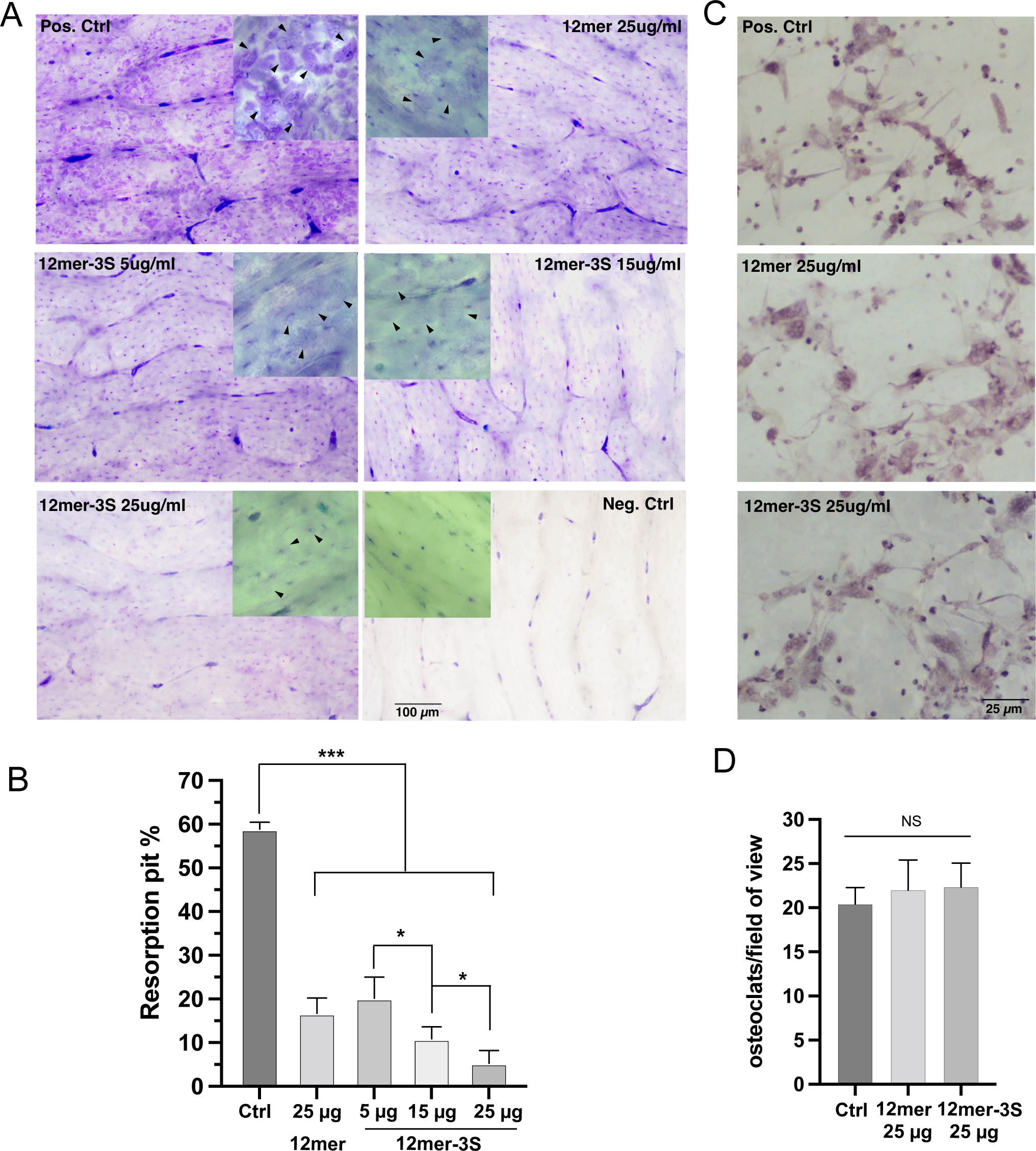
HS oligosaccharides inhibit bone resorption of murine osteoclasts. (A) Resorption of bovine bone slices by bone marrow-derived murine osteoclasts. After seeding with osteoclasts, the bone slices were either untreated (positive control),or treated with 25 µg/ml HS-12merNS2S6S (12mer) or 5, 15 and 25 µg/ml HS-12merNS2S3S6S (12mer-3S) for 6 days. The bone slices were stained with Toluidine blue to visualize the resorption pits. Images were taken with 10X lens. Of note, the bone slices show background staining between lamellar bone layers (see stained negative control bone slice), these strongly stained areas are not caused by osteoclasts resorption. Inset: images taken at 20X lens, and the black arrowhead indicate resorption pits generated by osteoblasts. (B) Quantification of percentage of bone surface area covered by resorption pits. n = 3. *** represents p < 0.001, * represents p < 0.05, by Student’s t test. (C) Separate sets of bone slices were seeded with murine osteoclasts and were either untreated, or treated with 25 µg/ml HS-12merNS2S6S or HS-12merNS2S3S6S for 6 days. The bone slices were processed by TRAP staining to virtualize multinucleated osteoclasts (cells stained purple) on the bone slices. Images were taken with 20X lens. (D) Quantification of the number of osteoclasts (defined as TRAP-positive cells with more than 3 nuclei) per 20X field of view. n = 4.

## Discussion

In this study, we sought to understand molecular details of the CtsK-HS interaction and how this interaction might alter the structure and function of CtsK. We show for the first time that HS oligosaccharides can induce monomeric CtsK to form a stable dimer. Crystal structures, combined with mutagenesis, revealed residues involved in HS interactions and suggest possible modes of dimerization. It is interesting to note that the asymmetric unit dimer (A/B), combined with the crystallographic symmetry-related dimer (A/A’) (Fig. 4A), can be repeated in a helical fashion along a theoretically extended HS chain maximizing the concentration of CtsK per HS chain at the cell surface (Figure S4). These structures also reveal how Cst3 binds to and inhibits CtsK by showing the specific interactions between the N-terminus of Cst3 and the active site of CtsK and that this binding neither interferes with CtsK dimerization nor HS binding.

The exact mechanism by which CtsK digests collagen fibrils remains poorly understood. Unlike other collagenases, such as matrix metalloproteases (MMPs), and bacterial collagenases, which all have additional collagen-binding domains that engage in binding and unwinding of the collagen triple helix (33–36), CtsK appears to be handicapped with only a single domain. To explain the collagenase activity of CtsK, the prevailing view in the field is that multiple molecules of CtsK likely organize into an oligomer on the surface of collagen fibril, with some molecules helping to unwind the triple helix while some perform the proteolysis (28). The exact form of this proposed oligomer remains a mystery. One study suggested that a crystallographic dimer in their crystal might function as a unit to unwind and digest collagen triple helix (28). In this model, the authors also proposed an essential role of CS, which is known to be present in the collagen fibril, in recruiting CtsK to the surface of the collagen fibril. Because most of this model is based on molecular docking of tropocollagen to the dimer, determining whether this model is truly physiologically relevant requires more in-depth study.

However, regardless of the exact mechanism by which CtsK digests collagen fibrils, our study revealed that CtsK-HS interaction strongly interferes with the digestion mechanism. We envision that HS might interfere with the collagenase activity of CtsK through at least two distinct mechanisms. If multiple molecules of CtsK are indeed required for activity, stable binding of HS to CtsK might directly interfere with the efficiency of assembling the correct CtsK oligomer with collagenase activity. If binding to CS is indeed required for initiating digestion, then HS could inhibit collagenase activity by directly suppressing the CtsK-CS interaction because the binding sites of CS and HS on CtsK partially overlap, and that HS is known to bind CtsK with higher affinity (17).

It’s important to note the molecular mechanism of CtsK inhibition by HS is fundamentally different from conventional means of protease inhibition. The most common way of protease inhibition is by targeting the active site with small molecules. This has been explored very successfully in blocking the activity of CtsK and has been tested in clinical trials (37, 38). The main problem of this type of inhibitors is that it indiscriminately inhibits both peptidase and collagenase activities of CtsK, which can interrupt normal peptidase activity of CtsK and potentially lead to side effects. A different approach would be to design inhibitors to target exosites of CtsK, with the goal of selectively inhibiting the collagenase activity of CtsK without inhibiting the peptidase activity (15). Several of this class of inhibitors have been described (16, 39), but there is still a long way to go in terms of demonstrating the efficacy and safety of these novel compounds. In contrast, the chief mechanism of inhibition by HS may be through changing the oligomeric state of CtsK and/or blocking collagenase binding. As we have demonstrated clearly, this mechanism inhibits collagenase activity only, which reduces the risk of side effects due to undesired inhibition of peptidase activity. Compared to newly developed exosite inhibitors, HS-based oligosaccharides bear a unique advantage in that their safety profiles are well documented due to extensive use of HS-based oligosaccharides as anti-coagulant drugs (40, 41). A third unique feature of HS-mediated inhibition of CtsK is that it inhibits without totally blocking the collagenase activity of CtsK. This at first glance might look like a disadvantage for an inhibitor, but in the arena of curbing bone resorption, this is likely a better choice than a complete collagenase blocker. Normal levels of bone resorption are required to maintain healthy bone structure by renewing bones that are worn done due to microfractures. If CtsK activity is completely shut down these microfractures will remain unrepaired. Long term, this could greatly increase the incidence of fracture. On the other hand, if CtsK activity is reduced to a lower level, then one can reduce excessive bone resorption while maintaining low level of activity to allow bone repair. Combined, HS-based oligosaccharides represent a brand-new strategy of inhibiting pathological bone resorption.

Our discoveries also raised many intriguing questions regarding how HS might regulate the physiological function of CtsK during bone resorption. Due to highly abundant expression of both CtsK and HS by osteoclasts (Fig.7), cell surface HS could capture newly synthesized CtsK, sequestering it at the cell surface until needed. Under this scenario, HS could serve as a buffer zone to regulate the amount of CtsK being released into bone matrix for active bone resorption. The cell surface HS-bound CtsK can either be kept active for extended periods for later release into resorption pit, or it can be internalized by membrane-attached HS proteoglycans (HSPGs) to reduce extracellular concentration of CtsK and slow down bone resorption. The interaction could also happen in the extracellular matrix when matrix HSPGs (such as perlecan and agrin) are secreted directly into the resorption pits, or when membrane-attached HSPGs (such as syndecans) are actively shed into the resorption pit. In this way osteoclasts might temporarily slow down bone resorption without altering overall expression levels of CtsK. Other possibility of interaction also exists, such as HS might be involved in autoactivation of pro-CtsK as it is also a HS-binding protein and they might encounter each other in the secretory vesicles and also in the extracellular space. We are actively investigating all these possibilities and our future studies will shed more light on the physiological roles of CtsK-HS interaction in bone resorption.

In conclusion, our study on CtsK-HS interaction has led to understanding of novel mechanisms that regulates the enzymatic activities of CtsK. These data not only shed new light on how CtsK functions in bone resorption, they will also be essential for understanding the role of CtsK in other pathophysiological conditions, such as cartilage degeneration, cardiovascular diseases and cancer. We plan to use this study as a foundation to design a new class of CtsK inhibitor, which can be further developed into an anti-resorptive therapeutics to curb undesired bone resorption in diseases like osteoporosis, periodontal bone loss, rheumatoid arthritis, Paget’s disease and cancer.

## Materials and Methods

### Expression and purification of full-length murine procathespin K (pro-CtsK) in mammalian cells

Complete open reading frame of murine pro-CtsK (GE Dharmacon) were cloned into pUNO1 (Invivogen) expression vector using AgeI and NheI restriction sites. The sequence of the final plasmid was confirmed by DNA sequencing. Recombinant murine pro-CtsK were expressed in 293-Freestyle cells (ThermoFisher) by transient expression using FectoPRO transfection reagent (Polyplus transfection). Purification of pro-CtsK from culture supernatant was carried out using HiTrap heparin-Sepharose column (GE Healthcare) at pH 7.1 (HEPES buffer). After purification, procathepsin K was >99% pure as judged by silver staining.

### Activation of pro-CtsK and purification of active CtsK

The conversion of the pro-CtsK into the active enzyme was accomplished by an autoactivation process. Briefly, the purified pro-CtsK was diluted with the same volume of the solution containing 0.1 M sodium acetate, 2.5mM EDTA, 10 mM DTT at pH 4. Mature CtsK seed was added into pro-CtsK at 1:200 mass ratio, and the mixture was incubated at 4°C for 3-4 days. The activation was monitored at UV 405nm by using a peptide substrate Z-Phe-Arg-pNA. Mature CtsK was purified from the autodigestion mixture by HiTrap heparin-Sepharose column upon completion of autoactivation. CtsK was eluted with a salt gradient from 150 mM to 2 M NaCl (in 20mM NaOAc, pH 5.5). Purified CtsK was >99% pure as judged by silver staining.

### Heparin–Sepharose chromatography

To characterize the binding of pro-CtsK and mature CtsK to heparin, ∼50 μg of purified WT or mutant CtsK was applied to a 1 ml HiTrap heparin– Sepharose column (Cytiva Lifesciences) and eluted with a salt gradient from 150mM to 2M NaCl at pH 5.5 in 20 mM NaOAc buffer. The conductivity measurements at the peak of the elution were converted to the concentration of NaCl based on a standard curve.

### Surface plasmon resonance

Biotinylated 12merNS2S6S (compound # 20) was immobilized onto a streptavidin sensor chip (Nicoya) and analyzed on an OpenSPR instrument (Nicoya). In brief, 150 µl of solution of the 12mer-Biotin conjugate (20 µg/ml) in HBS-running buffer (25 mM HEPES, pH 7.1, 150 mM NaCl, 0.05% Tween-20) was injected to channel 2 of the flow cell at a flow rate of 20 µl/min. Immobilization of HS was confirmed by an increase of 100-200 resonance unit in channel 2. The flow cell channel 1 without any immobilization was served as a background control. Different dilutions of CtsK (concentrations from 22 to 360nM) in HBS-running buffer were injected at a flow rate of 20 µl/min for 3 mins followed by a 2-minute dissociation phase. The sensor surface was regenerated by injecting with 150 µl regeneration buffer (25 mM HEPES, pH 7.1, 2M NaCl) at a flow rate of 150 µl/min. The sensorgrams were fit with 1:1 Langmuir binding model from TraceDrawer 1.9.2.

### Glycan array binding assay

HS oligosaccharide array with 52 structure-defined HS oligosaccharides was prepared as previously described (42). Briefly, the amine containing HS oligosaccharides were printed onto NHS-activated glass slides. Each compound was printed in a 6 X 6 pattern. The microarray was probed with 1 μg/ml biotinylated-CtsK in 25mM HEPES buffer containing 150mM NaCl at pH 7.1. Bound CtsK was stained with an avidin Alexaflour-488 conjugate. The images were acquired using GenePix 4300 A scanner. The intensity data is the mean value ± S.D. of 36 individual spots.

### Analytical size-exclusion chromatography (SEC)

For analysis of CtsK and HS oligosaccharide complexes, purified mature CtsK (40 µg) was incubated with HS oligosaccharides (molar ratio 1:1) in 20 mM NaOAc, 250 mM NaCl, pH 5.5 for 1h at room temperature. Complexes were resolved on a Superdex 200 Increase column (Cytiva Lifesciences) using 20 mM NaOAc, 250 mM NaCl, pH 5.5, at 4°C. Please note, the presence of a para-nitrolphenyl group on the reducing end of the oligosaccharides allows excess oligosaccharides to be visible in the A280 elution profile.

### SEC-Multiangle light scattering (MALS)

SEC-MALS analysis was performed using a DAWN MALS detector (Wyatt Technology) connected to an AKTA FPLC system (GE Healthsciences). CtsK, CtsK/12merNS2S6S (compound # 20) and CtsK/12merNS6S (compound # 24) complexes were prepared as described above and concentrated to ∼2mg/ml, and 100 µl was resolved on Superdex 200 Increase SEC column using 20 mM NaOAc, 250 mM NaCl, pH 5.5. The MALS data were analyzed using ASTRA software (ver. 7.3.2.17).

### Enzymatic assay of the peptidase activity of CtsK

CtsK activity was determined using either a fluorogenic substrate in a microtiter plate format or a colorimetric substrate in a cuvette method. For routine analysis of CtsK activity (such as during autoactivation process), the colorimetric substrate was used. The reaction mixture consists of 500 ng of mature cathepsin K, 40 µg z-Phe-Arg-pNA (Bachem) in 150 µl CtsK digestion buffer (100 mM sodium acetate, 5 mM DTT, 5 mM EDTA at pH 5.5). A405 measurements were taken in a spectrophotometer (BioMate 3, Thermo Fisher) for 5 min with 1 min intervals. To examine the inhibitory effects of HS, Cst3 inhibition of Ctsk, and the stability of CtsK, the fluorogenic substrate (z-Leu-Arg-AMC) was used. The reaction mixture consists of ∼200 ng mature cathepsin, 5 µg z-Leu-Arg-AMC in 150 µl CtsK digestion buffer. The fluorescence signal (excitation: 370 nm, emission:450 nm) was measured on a fluorescence plate reader (Molecular Devices) for 15 min with 1 min intervals.

For digestion of the extracellular domain of human RAGE (soluble RAGE, or sRAGE), 5 μg sRAGE was digested by 20 ng mCtsK in CtsK digestion buffer at 22 °C in the presence of absence of 12mer for up to 20 mins.

### Enzymatic assay of the collagenase activity of CtsK

For digestion of bovine type I collagen (Sigma), 5 μg collagen was digested by 100 ng CtsK in 25 µl CtsK digestion buffer in the presence or absence of 12mer (1:2 molar ratio) at 37 °C for 2 hours. The reaction mixture was resolved on a 4-20% Bis-Tris gel (Genscript) and visualized by Coomassie staining.

For kinetic analysis of collagenase activity, 10 μg bovine type I collagen-FITC conjugate (ThermoFisher) was digested by 500 ng CtsK in 200 µl CtsK digestion buffer in the presence or absence of different 12mers in various amounts as indicated in the figure legends (molar ratio Ctsk:12mer = 4:1 to 1:2.5) at 22 °C. The fluorescence signal (excitation: 485 nm, emission: 525 nm) was measured on a fluorescence plate reader (Molecular Devices) for 30 min with 1 min intervals.

### Stability assay of mCtsK

The stability of CtsK assessed by storing either CtsK alone, or CtsK together with heparin (4:1 molar ratio), in a buffer containing 2.5 mM MMTS (a reversible active site inhibitor of cysteine proteases) and 50 mM NaOAc (pH 5.5) at 4 °C for 7 days. On day 1, 2, 3, 5 and 7, ∼ 200 ng aliquots were taken to measure the peptidase activity using the fluorogenic substrate as describe above. Also on selected days, 40 µg of CtsK or CtsK/heparin complex were analyzed by heparin-Sepharose chromatography as described above.

### Crystallization of CtsK-12mer and Cst3-CtsK-12mer complexes

The CtsK-12mer complex was generated by mixing mature CtsK with 12mer-NS2S6S in 1:1 molar ratio in 150mM NaCl, 20mM NaOAc, pH5.5 at 4°C overnight. The complex was purified by G2000SW 21.5mm × 60cm column (Tosoh) using 20mM NaOAc, 150mM NaCl, pH5.5. The purified complex was concentrated using a 10-kDa molecular mass cutoff Centricon tube (Millipore) to a final concentration of 6 mg/ml for crystallization trials. Cst3-CtsK-12mer ternary complex was generated by mixing Cst3, CtsK and 12mer-NS2S3S6S in 2:1:1 molar ratio in 150mM NaCl, 20mM NaOAc, pH5.5 at 4°C overnight. The complex was purified by SEC as describe above and was concentrated to 6 mg/ml for crystallization trials.

The CtsK/12mer (compound #20) complex was crystallized at room temperature using the hanging drop vapor diffusion technique by mixing 1.5 µl of protein solution with 1.5 µl of the reservoir consisting of 0.1M sodium acetate pH 5.0, 200mM NaCl, 31% PEG8000. Crystals were harvested by adding 1ul of cryo solution, consisting of 85% reservoir solution and 15% ethylene glycol, directly to the drop. The crystal was then transferred to 100% cryo-solution twice, followed by flash freezing in liquid nitrogen for data collection. Crystals of the Ctsk/Cst3/12mer (compound #19) complex were also obtained at room temperature by mixing 400 nl of protein solution with 400nls of the reservoir consisting of 92.5mM sodium citrate pH 5.6, 10mM barium chloride, 185 mM ammonium sulfate and 23 % PEG4000 using the sitting drop vapor diffusion technique. For data collection, 1ul of cryo-solution consisting of 85mM sodium acetate, 2mM barium chloride, 185 mM ammonium sulfate, 21.25 % PEG4000 and 15% ethylene glycol was added directly to the drop, then transferred to 100% cryo-solution, followed by flash freezing in liquid nitrogen. Data were collected on the Southeast Regional Collaborative Access Team (SER-CAT) 22-ID beamline at the Advanced Photon Source, Argonne National Laboratory (Table S1). Data were integrated and scaled using HKL2000 (43). Structures were solved by performing molecular replacement using PDB coordinates 3C9E of human Ctsk for the Ctsk/12mer complex and coordinates 3C9E (29) and 3GAX of human cystatin C (44) for the Ctsk/Cst-3 complex with Phaser (45), followed by iterative cycles of refinement in Phenix and manual model building in Coot (46–49). Model statistics and quality were evaluated using MolProbity (50) and are presented in Table S1.

### Site-directed mutagenesis

CtsK mutants were prepared by site-directed mutagenesis and the mutations were confirmed by Sanger sequencing. Expression, purification and autoactivation of the mutants was carried out the same way as the WT CtsK. All mutants were expressed at a comparable level to WT CtsK and showed comparable autodigestion rate as WT CtsK.

### Fluorescence-Activated Cell Sorting

MC3T3-E1 cells were lifted from culture dish using Accutase (Biolegend) and incubated with WT or mutant CtsK at 100 ng/mL in PBS with 0.1% BSA for 1 hr at 4 °C. Bound CtsK was stained with our rabbit anti-mouse CtsK (1µg/mL) for 1 hr at 4 °C, followed by goat anti-rabbit IgG-Alexa 647 (1:1,000; ThermoFisher Scientific) for 30 min and analyzed by flow cytometry.

The rabbit anti-mouse CtsK antibody was developed by immunizing rabbit with a mixture of murine proCtsK and mature CtsK. The polyclonal antibody was affinity purified from the anti-serum with a column immobilized with proCtsK, and the purified antibody displayed strong binding to both proCtsK and mature CtsK.

### Immunohistochemistry

Animal use was approved by the Institutional Animal Care and Use Committee (IACUC) of the University at Buffalo (Buffalo, NY). Femurs from 10-week-old mice were harvested and fixed for 48 h in 10% neutral buffered formalin, and decalcified in 10% EDTA for at 22 °C for 14 days. The samples were embedded in paraffin and sectioned at 5 μm. After antigen retrieval with citric acid (pH 6) and blocking, sections were immuno-stained with 1 μg/ml rabbit anti-mouse CtsK polyclonal antibody (prepared as described above) or 1 µg/ml human anti-HS mAb (HS20) overnight at 4 °C. For IHC staining of CtsK, after washing with PBS the bone sections were treated for 1 h with biotinylated goat anti-rabbit IgG secondary antibody (1:200; Vector Laboratories). The sections were developed using the ABC system (Vector Laboratories) and the cell nuclei were counter stained with 15% Ehrlich’s hematoxylin (Electron Microscopy Sciences). IHC staining of HS were performed similarly except biotinylated rabbit anti-human secondary antibody was used. For immunofluorescence staining, slides were treated with anti-rabbit Alexa-594 to visualize CtsK and anti-human Alexa-488 to visualize HS. The nuclei were stained with Dapi.

The specificity of anti-CtsK polyclonal antibody was confirmed by an absence of CtsK staining in bone sections obtained from CtsK knockout mice. The specificity of mAb HS20 was confirmed by an absence of HS staining in heparin lyase III treated bone sections.

### Bone resorption assays and analyses

Bone marrow cells were isolated by flushing the femurs and tibias of 6-to 10-week-old C57BL/6 mice. To generate osteoclasts (OCs), the BMMs were seeded into 6-well plates and incubated with macrophage colony-stimulating factor (M-CSF) (20_ng/ml, Biolegend) and Receptor activator of nuclear factor kappa-Β ligand (RANKL) (50_ng/ml) in alpha-MEM supplemented with 10% FBS for 4–5 days. When multinucleated OCs start to appear, the OCs were detached by Accutase and seeded on bovine bone slices (0.3mm thickness, prepared from bovine femur cortical bone by low-speed diamond saw) (51) at a density of 15000 cells per bone slice, and cultured in the presence of M-CSF (10 ng/ml) and RANKL (50 ng/ml). Selected bone slices were treated with HS 12merNS2S6S (compound #20) or 12merNS2S3S6S (compound #19) at various concentration (5 to 25 µg/ml) with medium change every 2 days. After 6 days, bone slices were harvested and the resorption pit area was visualized by 1% toluidine blue staining. For each bone slice, the number of osteoclasts per field of view was averaged from 6 images taken with 10X objective lens. The percentage of bone surface covered by resorption pits was quantified using Image J by setting a threshold that best distinguish the blue resorption pit and the unstained bone background in the positive control sample. For each experiment, 4 bone slices were used for each condition. The experiment was repeated three times with similar results.

Separate sets of bone slices were seeded with murine osteoclasts and were either untreated or treated with 25 µg/ml HS-12merNS2S6S or HS-12merNS2S3S6S for 6 days as describe above. The bone slices were fixed and stained with a TRAP staining kit (Sigma-Aldrich) to virtualize multinucleated osteoclasts on the bone slices. These cells contain more than 3 nuclei and are stained purple with irregular shapes. For each bone slice (n = 4 for each condition), the number of osteoclasts per field of view was averaged from 8 images taken with 20X objective lens. The experiment was repeated twice with similar results.

### Statistical analysis

All data are expressed as means ± SDs. Statistical significance was assessed using two-tailed Student’s t-tests or analysis of variance (ANOVA) using GraphPad Prism software (GraphPad Sofware Inc.). p value <0.05 was considered significant.

## Author contributions

Xiaoxiao Zhang, Conceptualization, Formal analysis, Investigation, Methodology, Writing – original draft; Yin Luo, Huanmeng Hao, Juno M. Krahn, Guowei Su and Robert Dutcher, Formal analysis, Investigation, Methodology; Y.X. completed the synthesis of 12-mers; Jian Liu, Conceptualization, Funding acquisition, Resources, Supervision, Writing – review and editing; Lars Pedersen, Conceptualization, Formal analysis, Investigation, Methodology, Funding acquisition, Resources, Writing – original draft, review and editing; Ding Xu, Conceptualization, Data curation, Formal analysis, Funding acquisition, Investigation, Methodology, Project administration, Resources, Supervision, Validation, Writing – original draft, review and editing.

## Acknowledgments

Data were collected at Southeast Regional Collaborative Access Team (SER-CAT) 22-ID beamline at the Advanced Photon Source, Argonne National Laboratory. SER-CAT is supported by its member institutions, and equipment grants (S10_RR25528, S10_RR028976 and S10_OD027000) from the National Institutes of Health. Use of the Advanced Photon Source was supported by the U. S. Department of Energy, Office of Science, Office of Basic Energy Sciences, under Contract No. W-31-109-Eng-38., supported by DOE Office of Biological and Environmental Research. We thank Dr. Mitchell Ho (NCI) for generously gifting us anti-HS mAb HS-20. The authors thank Andrea Kaminski and Percy Tumbale for critical reading of the manuscript.

