## Supplemental Fig. S1-4, Table S1 for "Heparan sulfate selectively inhibits the collagenase activity of cathepsin K"

### Figure S1

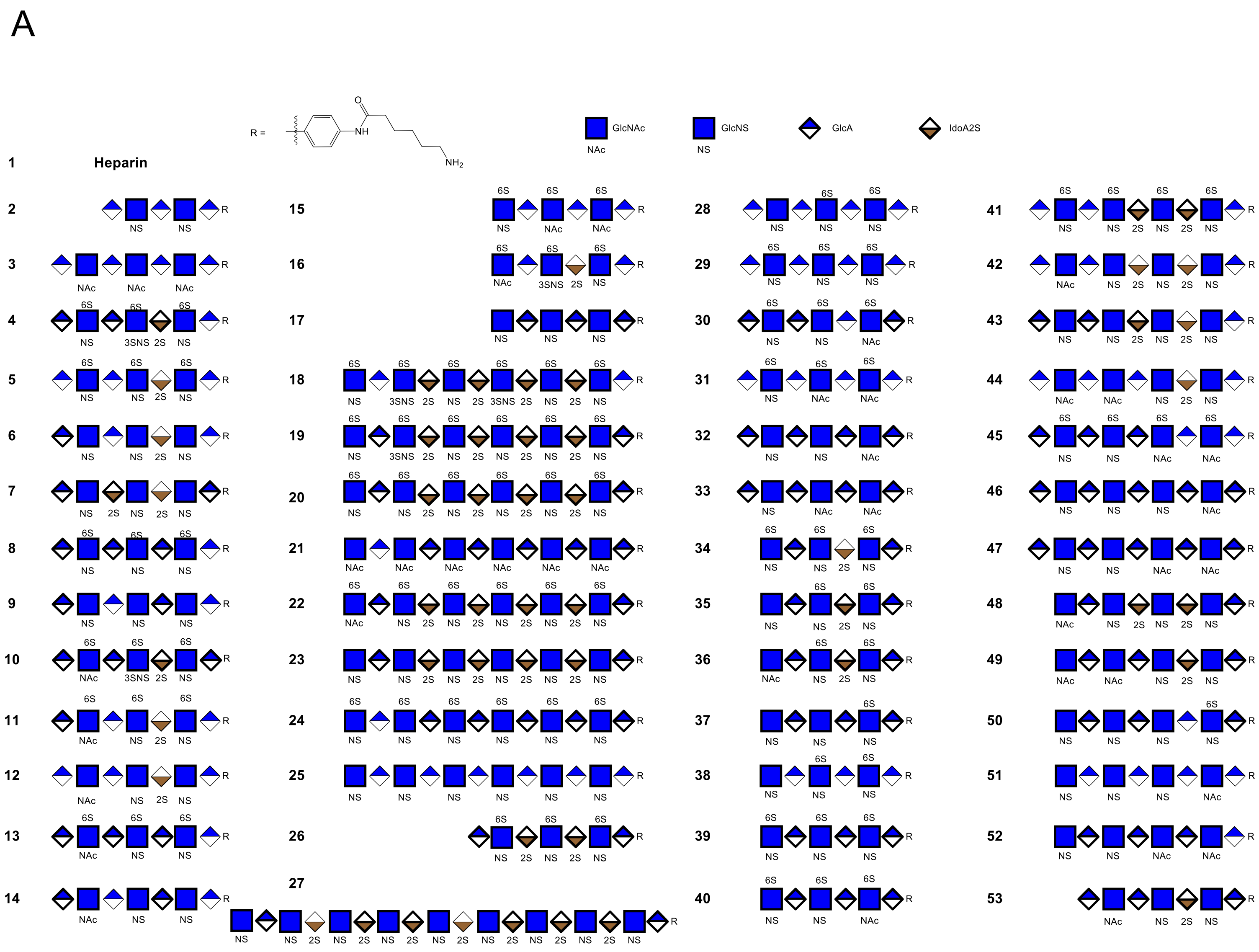

**Figure S1. Structures of HS oligosaccharides immobilized on the microarray.**

### Figure S2

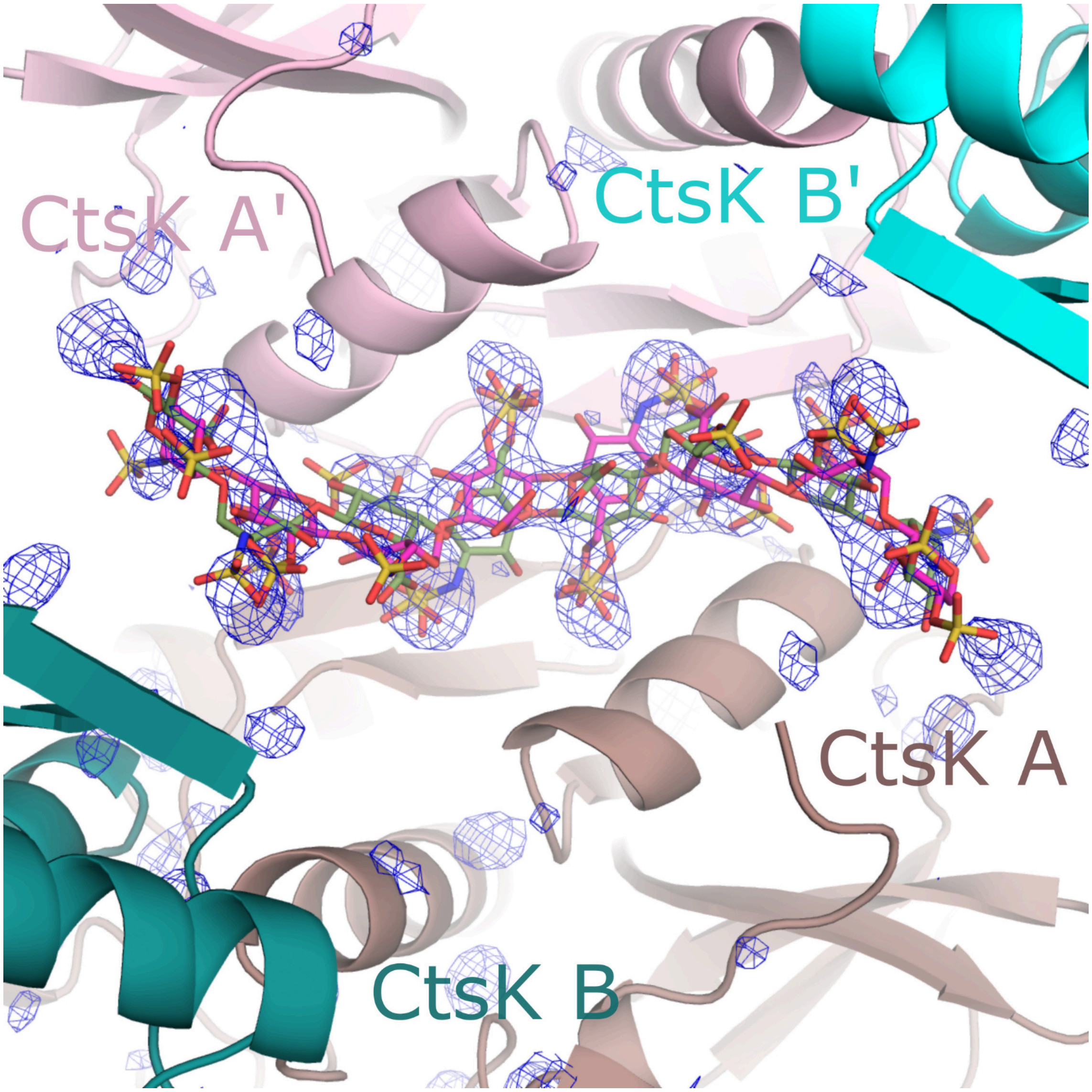

**Figure S2. Electron density of Heparan sulfate bound to CtsK.** Simulated annealing fo-fc omit density for the heparan sulfate oligomer bound in the Ctsk-12merNS2S6S structure is displayed contoured at 3σ. Symmetry related HS 12mers are shown in magenta and light green, respectively.

### Figure S3

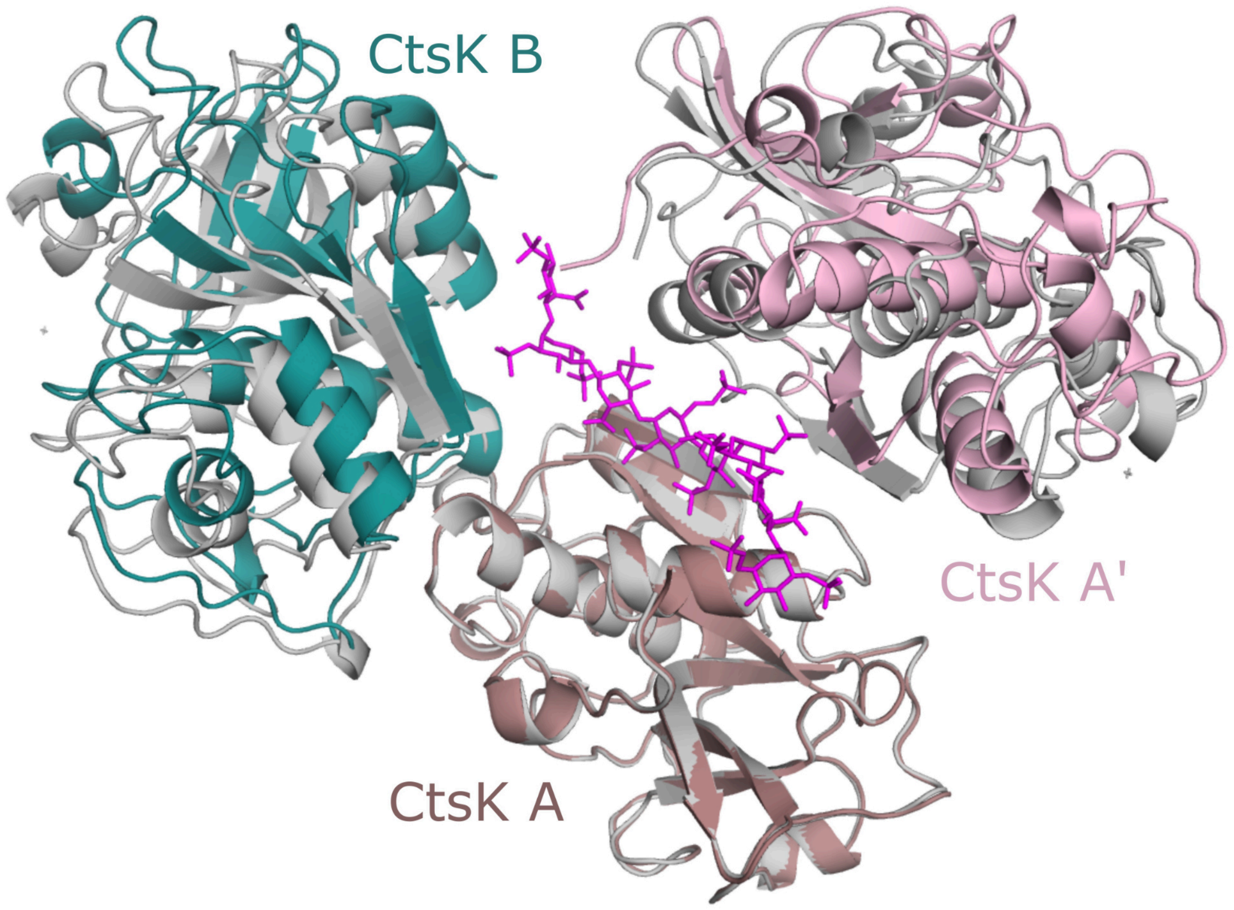

**Figure S3. Similar interfaces in the CtsK+12merNS2S6S and the CtsK+Cst-3+12merNS2S3S6S structures.** Superposition of monomer A from the crystallographic tetramer of Ctsk+12merNS2S6S (all molecules in grey, oligosaccharide in magenta) onto monomer A of the crystallographic tetramer found in the CtsK+Cst-3+12merNS2S3S6S present structure (monomers colored as in Figure 5). The symmetry related B' molecules from both structures and the Cst-3s have been removed for easy of viewing. Similar crystallographic interfaces exist in both structures.

### Figure S4

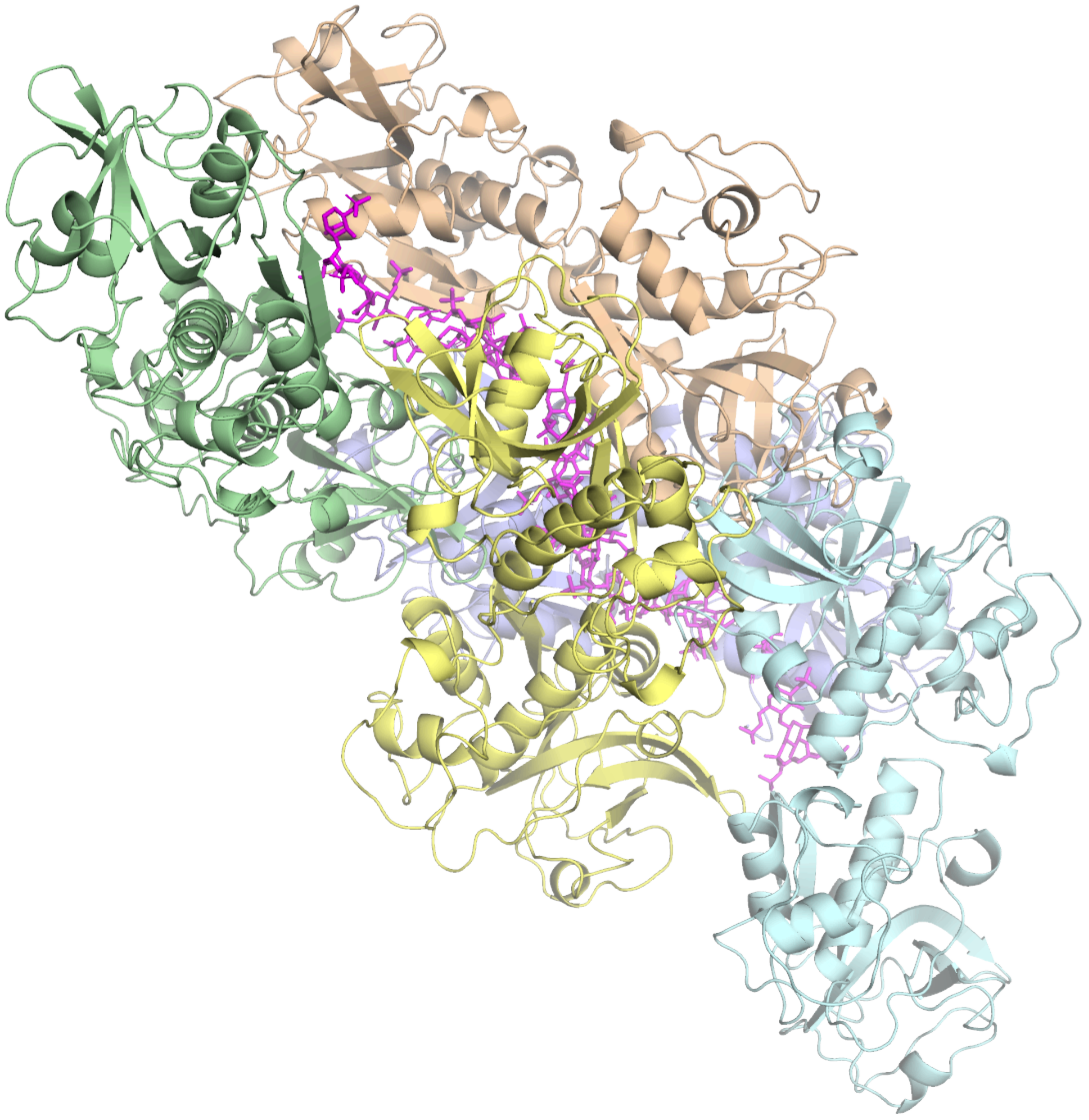

**Figure S4. Theoretical helical packing of CtsK along HS.** A theoretical model of CtsK binding to HS can be generated without much overlap by repeating the AB dimers (shown in pale cyan, light blue, light yellow, wheat and light green) with the same positioning as found at the crystallographic interface of the AA' dimer. The heparan is positioned based on the superposition of the CtsK. This model would allow for efficient packing of CtsK along the heparan sulfate chain.

**Table S1**. Crystallographic data and refinement statistics

| | CstK + 12merNS2S6S | CstK + CycC + HS[a] |
|---|---|---|
| PDB ID code | 8V58 | 8V57 |
| **Data collection** | 201116_SB3PN9 | 201009_SB1PN12 |
| Space group | $P4_122$ | $P222_1$ |
| Cell dimensions | | |
| *a*, *b*, *c* (Å) | 111.892, 111.982, 116.653 | 92.93, 179.61, 73.325 |
| α, β, γ (°) | 90, 90, 90 | 90, 90, 90 |
| Resolution (Å) | 50/0-3.1 (3.15-3.1)[b] | 50.00-2.75 (2.80-2.75) |
| $R_{sym}$ (%) | 9.2 (96.3) | 5.9 (77.0) |
| $I$ / σ$I$ | 5.4 (1.67) | 7.1 (1.66) |
| Completeness (%) | 98.8 (99.6) | 92.6 (86.6) |
| Redundancy | 7.4 | 7.3 (7.1) |
| **Refinement** | | |
| Resolution (Å) | 19.87-3.10 | 38.35-2.75 |
| No. reflections | 13,647 | 30,119 |
| $R_{work}$ / $R_{free}$ (%) | 23.59/29.26 | 22.86/25.79 |
| No. atoms | | |
| Protein | 3,169 | 9,813 |
| Water | 1 | 13 |
| Other | 252 | 115 |
| *B*-factors | | |
| Protein | 113.75 | 83.15 |
| Water | 99.06 | 69.18 |
| Other | 113.46 | 95.71 |
| R.m.s. deviations | | |
| Bond lengths (Å) | 0.009 | 0.007 |
| Bond angles (°) | 1.298 | 0.723 |
| Ramachandran Plot[c] | | |
| Allowed (%) | 4.93 | 5.22 |
| Favored (%) | 95.07 | 94.32 |

a) 12merNS2S3S6S was included in the crystallization but is not modeled in the structure due to the density being ambiguous.
b) High resolution shell is shown in parentheses.
